# Auxin antagonism of ABA-induced stomatal closure in *Arabidopsis* requires the TIR1/AFB–AUX/IAA module and ROS suppression

**DOI:** 10.1101/2025.07.15.664744

**Authors:** Camila Alejandra Rios, Laila Toum, Lucía Padula, Fernando Pieckenstain, Gustavo E. Gudesblat

## Abstract

Auxin inhibits ABA-induced stomatal closure, yet the underlying mechanism remains unclear. In this work we quantified 3-Indoleacetic acid (IAA) inhibition of ABA-triggered closure in *Arabidopsis thaliana* mutants and transgenic plants and observed that IAA inhibition of ABA-induced stomatal closure was lost or weakened in *tir1-1* and *axr3-1* mutants, affected in auxin perception. The same was found in *aux1-7, yuc5/8/9* and *pils5-2/2-5* mutants, and in YUCCA8 and PILS5 overexpressors, all altered in auxin synthesis or transport. Stomatal sensitivity to IAA was also reduced in *phyB-9* and *rbohd, toc1* and *cca1/lhy* mutants, in certain ecotypes, and in response to mild drought. H^+^-ATPase mutant *aha2-5* showed reduced stomatal sensitivity to IAA at 22°C but not at 35°C, while high-temperature-induced stomatal opening was not affected in *tir1-1*. IAA also caused partial degradation of the DII-VENUS auxin reporter, and inhibited ABA-induced ROS production in guard cells. Our results show that IAA counteracts ABA signalling via the TIR1/AFB–AUX/IAA module and ROS suppression, with genetic factors and mild drought modulating the response.

## Introduction

Stomatal pores change their aperture in response to a number of exogenous and endogenous cues to regulate gas exchange. Such regulation is achieved through changes in guard cell turgor, brought about by ion channel regulation and starch degradation. H^+^ pumping by AUTO-INHIBITED H^+^-ATPases (AHA), induced by stimuli such as blue light, causes proton extrusion, hyperpolarising the guard cell membrane. The more negative membrane potential opens inward-rectifying K^+^ channels, allowing K^+^ influx, accompanied by Cl^-^ or NO_3_-uptake via H^+^-coupled cotransport, and biosynthesis of malate as a counteranion. The resulting osmotic influx of water increases guard cell turgor, opening the pore (Li et al., 2022). Abscisic acid (ABA) is a stress hormone that binds cytosolic PYR/PYL receptors. This blocks clade-A PP2C phosphatases, which relieves inhibition of SnRK2/OST1 that, together with Ca^2+^-dependent CPKs and CIPKs, jointly phosphorylate plasma-membrane NADPH oxidases (RBOHD/F) and the S-and R-type anion channels SLAC1 and ALMT12, while also priming the outward K^+^ channel GORK. Anion efflux depolarises the guard-cell membrane, and in parallel ABA causes the inhibition of plasma-membrane H^+^-ATPases, causing cytosolic alkalinisation and preventing membrane repolarisation. The coordinated efflux of Cl^−^, NO3^−^, malate^2−^ and K^+^ rapidly diminishes guard-cell turgor and closes stomata within minutes, while ABA-driven transcriptional reprogramming secures sustained closure under prolonged drought (Kollist et al., 2014; Hsu et al., 2021; Waadt et al., 2022; Seller & Schroeder, 2023). One of the endogenous stimuli that favour stomatal opening is auxin, 3-Indoleacetic acid (IAA). This hormone, as well as synthetic auxins, can promote stomatal opening in the dark (Levitt et al., 1987; Dunleavy & Ladley, 1995), and inhibit ABA and CO_2_-induced stomatal closure at physiological concentrations (Snaith & Mansfield, 1982; Tanaka, 2006; Cho et al., 2012). However, species, light quality and compound-dependent variations have been reported (Pospíšilová, 2003). Auxins can be autonomously perceived by guard cells, where they can provoke changes in inward-rectifying K^+^ (Blatt & Thiel, 1994) and anionic (Marten et al., 1991) currents in a dose-dependent manner in *Vicia fava*. They can also induce cytosolic acidification and Ca^2+^ elevation (Irving et al., 1992) and activate H^+^ extrusion (Lohse & Hedrich, 1992) in guard cells. Also, auxin can reduce dark-induced nitric oxide (NO) (Xiao-Ping & Xi-Gui, 2006) and reactive oxygen species (ROS) (Song et al., 2006) production in guard cells.

Auxin is perceived by the TRANSPORT INHIBITOR RESPONSE 1/AUXIN SIGNALING F-BOX - AUXIN/INDOLEACETIC ACID (TIR1/AFB–AUX/IAA) module. Upon auxin perception, TIR1/AFB F-box receptors interact with Aux/IAA repressor proteins and cause their ubiquitination and degradation, thereby releasing AUXIN RESPONSE FACTOR (ARF) transcription factors, which then activate auxin-responsive genes. TIR1/AFB-dependent gene expression mediates different auxin responses to developmental, mechanical and environmental cues. (Leyser, 2018; Ang & Østergaard, 2023). Some TIR1/AFB-dependent responses can be very fast, such as the rapid auxin-induced apoplast acidification in *Arabidopsis thaliana* (Arabidopsis) hypocotyls, which stimulate cell elongation within minutes through the induction SAUR (Small Auxin Up RNA) genes, causing the activation of PM H^+^-ATPases (Fendrych et al., 2016).

Recently, it was found that TIR1/AFBs possess adenylate cyclase activity, which leads to the rapid production of cyclic adenosine monophosphate (cAMP), a second messenger required for TIR1/AFB–AUX/IAA-dependent gene expression (Chen et al., 2025). Another cyclic nucleotide, cyclic guanosine monophosphate (cGMP), is also synthesised by TIR1/AFBs through a guanylate cyclase motif independent from its adenylate cyclase motif (Qi et al., 2023).

In addition to the TIR1/AFB–AUX/IAA module, extracellular Auxin Binding Protein 1 (ABP1) and related ABP1-LIKE proteins can bind auxin with high affinity and transduce their signals through TRANSMEMBRANE KINASE (TMK) receptor-like kinases (Vanneste et al., 2025). TMKs can interact with, and rapidly activate, several plasma membrane AHAs by direct phosphorylation (Li et al., 2021; Lin et al., 2021). Two of them, AHA1 and AHA2, are highly expressed in Arabidopsis guard cell protoplasts (Ueno et al., 2005). While AHA1 plays a predominant role in stomatal opening in response to blue light (Yamauchi et al., 2016), AHA2 mediates responses to other stimuli. The a*ha2* mutant, but not *aha1*, was found to be defective in high-temperature-induced stomatal opening (Kostaki et al., 2020). However, a recent study found that the high-temperature-associated kinase TARGET OF TEMPERATURE 3 directly controls the activity of AHA1 to induce stomatal opening (Xu et al., 2025). The regulation of AHA2 activity by phosphorylation is complex, as it can either activate or inhibit its activity depending on the target residue. Auxin-induced SMALL AUXIN UP RNA (SAUR) proteins inhibit PP2C D clade phosphatases, which dephosphorylate Thr-947 of AHA2 in guard cells and causes its inactivation (Wong et al., 2020), while PSK5 inactivates it through Ser-931 phosphorylation (Fuglsang et al., 2007). AHA2 appears to be involved not only in stomatal opening but also in closure, as it was shown that ABA-activated BRI1-ASSOCIATED RECEPTOR KINASE 1 (BAK1) phosphorylated it at Ser-944, leading to its activation and contributing to ABA-induced cytoplasmic alkalinization and stomatal closure (Pei et al., 2022). Other proton transporters might also mediate the effect of auxin on guard cells, since mutants defective in the vacuolar Ca^2+^/H^+^ exchangers CAX1 and CAX3 are affected in IAA-induced hyperpolarization of the plasma membrane, and are also insensitive to IAA’s inhibitory effect on ABA-induced stomatal closure, an effect also observed through the application of the pH-dependent AUX1 auxin transporter inhibitor 2-naphthoxyacetic acid (Cho et al., 2012). Different lines of evidence show that the circadian clock also regulates stomatal H^+^-ATPase activity and long-term water use efficiency (WUE) (Simon et al., 2020; Kamrani et al., 2022). Clock components like the central regulator TIMING OF CAB EXPRESSION 1 (TOC1), LATE ELONGATED HYPOCOTYL (LHY) and CIRCADIAN CLOCK ASSOCIATED1 (CCA1) also offer points of integration of light and temperature cues (Casal & Qüesta, 2018).

Auxin synthesis is controlled mainly through the expression of multiple TRYPTOPHAN AMINOTRANSFERASE OF ARABIDOPSIS 1 (TAA1) and TAA1-RELATED transaminases and YUCCA

(YUC) flavin-containing monooxygenases, whose genes display remarkably tissue-specific expression, consistently with its critical role in plant growth and development. The most important regulatory level of auxin action is its spatiotemporal distribution across tissues and organs, achieved mainly through polar cell-to-cell auxin transport, which is regulated by endogenous and environmental cues (Yu et al., 2022; Vanneste et al., 2025). A complex network of subcellularly compartmentalised biosynthesis, conjugation and degradation processes regulates auxin metabolism. Auxin can cross the plasma membrane by passive diffusion into the cell and by active transport via transporters including the previously mentioned AUX1, while polarly localized PIN-FORMED (PIN) transporters export auxin unidirectionally into the apoplast. Endoplasmic reticulum-localized PIN and PIN-LIKES (PILS) proteins possibly mediate bidirectional transport of auxin across the endoplasmic reticulum membrane and likely play a role in their metabolism. PILS proteins increase cellular auxin accumulation but decrease auxin signalling, while *pils2, pils5* and *pils2/pils5* mutant Arabidopsis seedlings showed significantly higher free IAA levels (Barbez et al., 2012).

Auxin has emerged over the past decade as an important player in adaptation to water deficit conditions, acting in a concerted manner with other stress hormones such as ABA an ethylene (Blakeslee et al., 2019; Leftley et al., 2021; Abuzeineh & Abualia, 2023). Several drought-induced DREB/CBF transcription factors induce the expression of IAA5 and IAA19, which belong to the family of Aux/IAA auxin-sensitive repressors, and whose mutation results in decreased tolerance to stress conditions (Shani et al., 2017). Further studies showed that IAA5, IAA6, and IAA19 act in a transcriptional cascade seemingly ABA-independent that maintains expression of glucosinolates when plants are exposed to drought. These secondary metabolites can trigger stomatal closure, and consistently with these observations, different multiple *tir1/afb* auxin receptor mutants are more resistant to mild drought (Salehin et al., 2019). In guard cell protoplasts, IAA levels rapidly decrease after ABA treatment (Jin et al., 2013; Zhu & Assmann, 2017).

Phytochrome B (PHYB), in combination with other receptors, can perceive light and temperature and modulate plant growth and development to improve adaptation to shade (low red:far red ratio) and high temperature. Both stimuli cause rapid auxin biosynthesis through TAA1 and YUC enzymes (Casal & Qüesta, 2018; Legris et al., 2019), and photoactivate d PHYB was shown to interact directly with AUX/IAA proteins to inhibit auxin signalling in Arabidopsis (Xu et al., 2018). As a result, *phyB* mutants display an elongated phenotype. Because PHYB normally represses auxin biosynthesis and transport genes under red light, in *phyB* these genes are upregulated, leading to increased auxin accumulation and transport, which drives exaggerated elongation growth.

It is well known that plants subjected to a prior dehydration/recovery cycle show a decreased rate of water loss in a subsequent dehydration stress, a phenomenon known as stress memory (Ding et al., 2012). Upon rehydration, stressed plants display reduced stomatal aperture for a period extending up to several days, something that has been linked to changes in guard cell gene expression in Arabidopsis (Virlouvet & Fromm, 2015; Seller & Schroeder, 2023) and rice (Auler et al., 2021). Unlike elevated CO_2_, a stimulus that also triggers stomatal closure, ABA treatment triggers extensive and dynamic chromatin remodelling in guard cells, with distinct changes in chromatin accessibility that persist for at least 24 h (Auler et al., 2021; Seller & Schroeder, 2023). Chromatin remodelling can be modulated by post-transcriptional gene silencing (PTGS) (Tresas et al., 2025), which has not been much studied in guard cells.

Many natural Arabidopsis accessions are widely distributed in various geographical areas. These diverse natural habitats have exerted different evolutionary pressures, which have allowed the accessions to develop different molecular and physiological strategies to adapt to their specific environment (Alonso-Blanco et al., 2016). Several reports have shown differences in adaptation to drought, as well as variation in stomatal response (Bouchabke et al., 2008; Vile et al., 2012; Aliniaeifard & Van Meeteren, 2014; Monroe et al., 2018; Chen et al., 2021). Therefore, understanding the genetic basis of this natural variation might provide clues on how plants can adapt to different precipitation regimes.

Although the effect of auxin on stomatal aperture is well documented in the scientific literature, it is not very clear how they are perceived, or which factors can affect the sensitivity to them. In this work, we found that a functional TIR1/AFB–AUX/IAA module is required for auxin inhibition of ABA-induced stomatal closure in Arabidopsis. Additionally, we found that stomatal sensitivity to auxin is reduced in different mutants or transgenic lines affected in auxin synthesis, transport, circadian rhythm, ROS production and in *aha2-5* mutant. The capacity of auxin to inhibit ABA-induced stomatal closure was also significantly reduced by mild drought, and in some Arabidopsis natural accessions. These results provide novel insights into auxin perception in guard cells.

## Materials and methods

### Plant material and growth conditions

Seeds were imbibed in water and stratified for 3 days in darkness at 4°C, then sown in either TS1 (Klasmann-Deilmann, Germany) or Growmix Multipro (Terrafertil, Argentina), two peat based substrates, mixed in both cases with sand in a 2:1 proportion (substrate:sand). Plants were grown in growth chambers under a 12h:12h light/dark cycle (photon flux density of 90 μE) at 22°C to 23°C. After a week, seedlings were transferred to larger pots and watered three times a week to 65% field capacity, by weighing individual pots. For drought treatments, well watered plants were watered as described, maintaining them between 50% and 65% field capacity, while plants under water deficit were maintained on a similar watering regime until day 20, when they were left unwatered until they reached 30 % field capacity (typically after five days), then watered to 40% field capacity. Somatal aperture experiments were performed two days after rewatering. Experiments were performed with *Arabidopsis thaliana* Columbia-0 (Col-0) ecotype and the following mutants and transgenic lines, all in Col-0 background: *tir1-1* (Ruegger et al., 1998), *axr3-1 (Leyser et al*., *1996), rbohD* (Torres et al., 2002), *aha2-5* (Haruta et al., 2010), *toc1* (Strayer et al., 2000), *cca1/lhy* (Alabadí et al., 2002), *aux1-7* (Pickett et al., 1990), *pils5-2/2-2* (Barbez et al., 2012), *phyB-9* (Reed et al., 1993) and triple YUCCA mutant *yuc5/yuc8/yuc9* (Chen et al., 2014), YUCCA8 OE (Hentrich et al., 2013), 35S:PILS5 (Barbez et al., 2012), *ago4-1* (Zilberman et al., 2004), DNA demethylase triple mutant *rdd* (*ros1/dml2/dml3*) (Le et al., 2014), *rdr2* (Xie et al., 2004), *rdr6* (Peragine et al., 2004), *dcl2/3/4* (Henderson et al., 2006) and DII-VENUS (Brunoud et al., 2012). For natural variation experiments, the following ecotypes were used: Li-7, Wei-0, St-0, Cvi-0, Mt-0 and An-1.

### Chemicals

3-Indoleacetic acid (IAA), 1-naphthaleneacetic acid (NAA), 2,4-dichlorophenoxyacetic acid (2,4D) were dissolved in DMSO, while ABA (mixed isomers) was solubilised in ethanol and coronatine (COR) in methanol. All reagents were purchased from PhytoTech Labs (USA).

### Stomatal aperture bioassays

Stomatal bioassays were performed essentially as previously described (Gudesblat et al., 2009). Pieces from the younger, fully developed leaves from 3-4-week-old plants were floated on 10 : 10 buffer (10 mM KCl and 10 mM MES-KOH, pH 6.15) under light for at least 2 h with the abaxial side in contact with the solution. Then IAA, NAA or 2,4 D (at indicated concentrations) were applied 10 minutes prior to addition of ABA (20 μM). 1 h after ABA addition, epidermis were obtained with double-sided tape and observed under the microscope. Stomatal apertures were measured using Image J 1.54 Software. For mock treatments, no compounds were added. High temperature assays in guard cells were performed as previously described (Kostaki et al., 2020), but incubating leaf pieces instead of epidermal peels. Leaf pieces were floated on 10/50 buffer (10 mM MES and 50 mM KCl, pH 6.15) at 22°C for 2 h, then transferred to fresh 10/50 buffer at 22 °C or prewarmed to 35°C and incubated for a further 2 h.

### Confocal microscopy

DII-VENUS abaxial epidermal peels from 3-4-week-old plant leaves were floated on 10:10 buffer (10 mM KCl and 10 mM MES-KOH, pH 6.15) under light for 2.5 h, then IAA (10 μM) or COR (1,56 μM) were added to the incubation medium. Peels were incubated for 60 minutes and observed in a confocal laser-scanning microscope (Carl Zeiss LSM 980, 405-nm excitation with argon laser line, and 488-nm long-pass emission) equipped with a Zeiss Plan-Apochromat 20x/0.8 M27 objective lens. Fluorescence in guard cell nuclei was analysed using ImageJ 1.54 software.

### ROS measurements

Hydrogen peroxide production in guard cells was measured using H_2_DCFDA (Murata et al., 2001). After a 2.5 h incubation in 10:10 buffer under light conditions, epidermal peels from 3-4-week-old plant leaves were transferred to 10 mM Tris-HCl pH 7.2 buffer containing H2DCFDA (10 μM) for 15 min. Excess H2DCFDA was removed by washing three times with 10 mM Tris-HCl pH 7.2. Then peels were transferred to 10:10 buffer containing the indicated treatments and incubated for 20 min in the dark. IAA (10 μM) was added 10 min prior to ABA (20 μM). Fluorescence was observed with a Nikon Eclipse E600 fluorescence microscope (excitation 460–480 nm, emission 495–540 nm). The fluorescence in guard cells was analysed using ImageJ 1.54 software.

### Statistical analysis

Statistical differences were determined by ANOVA (R package v 4.5.0) and Tukey′s test for pairwise comparisons. Before conducting the ANOVA test, DHARMa and glmmTMB packages in R were used to assess the assumptions of homoscedasticity and normality of residuals. glmmTMB package (Brooks et al., 2017) was employed to fit generalised linear mixed models with treatments (hormones or temperature) and genotype as explanatory variables, and the replicates as a random variable. The variance was modelled with dispformula. (Hartig, 2024) provided diagnostic tools to simulate and visually inspect residuals. The graphics were made with the ggplot2 package (Wickham, 2016).

## Results

It was previously shown that the naturally occurring auxin IAA can antagonise ABA-induced stomatal closure in Arabidopsis (Snaith & Mansfield, 1982; Tanaka, 2006; Cho et al., 2012). We performed dose-response curves with IAA as well as with synthetic auxins 1-naphthaleneacetic acid (NAA) and 2,4-dichlorophenoxyacetic acid (2,4-D). In all three cases we observed that auxins inhibited stomatal closure induced by ABA in a dose-dependent manner in the 0.1 to 10 μM range (Fig. 1).

**Figure 1.**
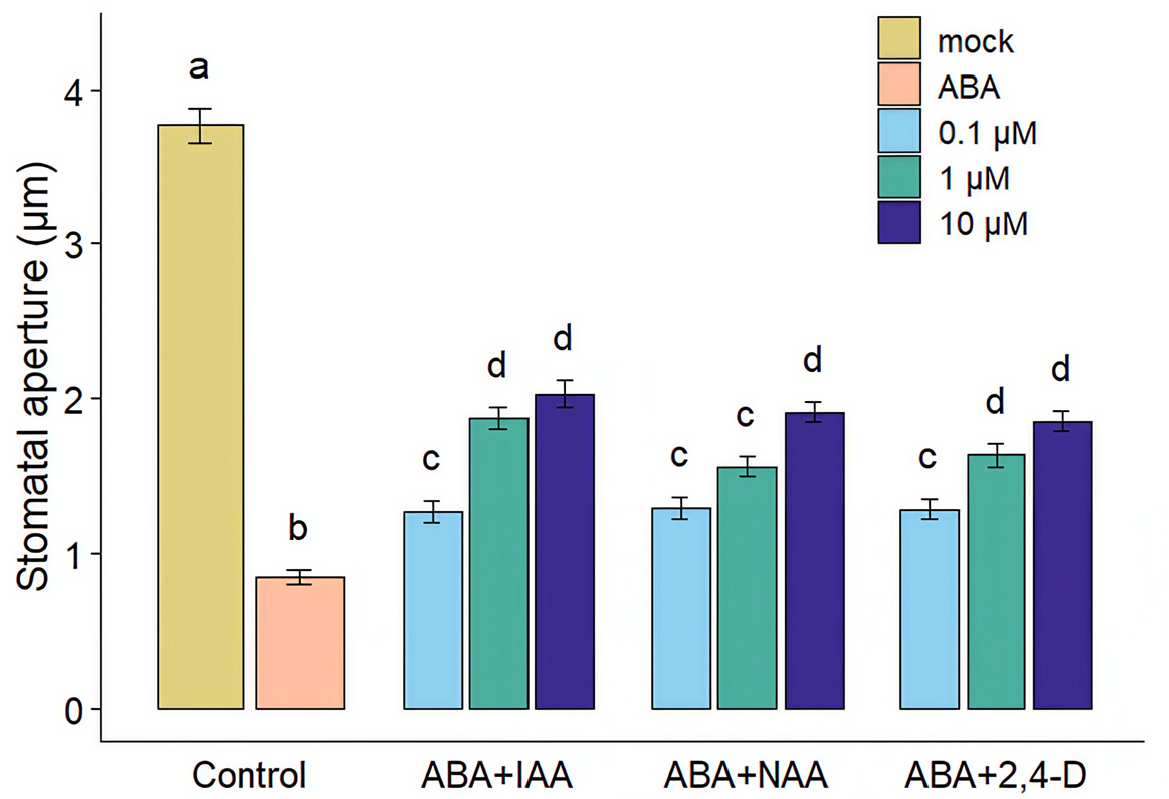
Auxins reduce ABA-induced stomatal closure in a dose-dependent manner. IAA, NAA and 2,4 D (at 0,1, 1 and 10 μM) were added 10 min prior to ABA (20 μM), and stomatal apertures were measured 1 h after ABA treatment. Different letters indicate significant differences at p < 0.05 (one-way ANOVA, Tukey′s test). Error bars represent SE from three independent trials, n=40 per trial in all experiments.

Next, we studied if inhibition of ABA-induced stomatal closure by IAA involves the TIR1/AFB–AUX/IAA signalling module. For this purpose, we tested *tir1-1* and *axr3-1* mutants. The first one lacks one of the auxin receptors, while the second one carries a dominant negative mutation in IAA17 repressor (Leyser et al., 1996). Both displayed stomatal insensitivity to IAA (Fig. 2a,b), indicating that the TIR1/AFB–AUX/IAA module is required for auxin inhibition of ABA-induced stomatal closure. In addition, we found evidence that TIR1/AFB–AUX/IAA is functional in Arabidopsis guard cells since IAA caused partial degradation in guard cells of the auxin perception reporter 35S:DII-VENUS, in which the N-terminal degron domain of IAA28 is fused to the VENUS fluorescent protein Fig. 2c) (Brunoud et al., 2012). No changes in DII-VENUS fluorescence were observed when peels were treated with the bacterial toxin coronatine, produced by *Pseudomonas syringae* pv. t*omato*, which is also capable of inhibiting ABA-induced stomatal closure (Melotto et al., 2006)

**Figure 2.**
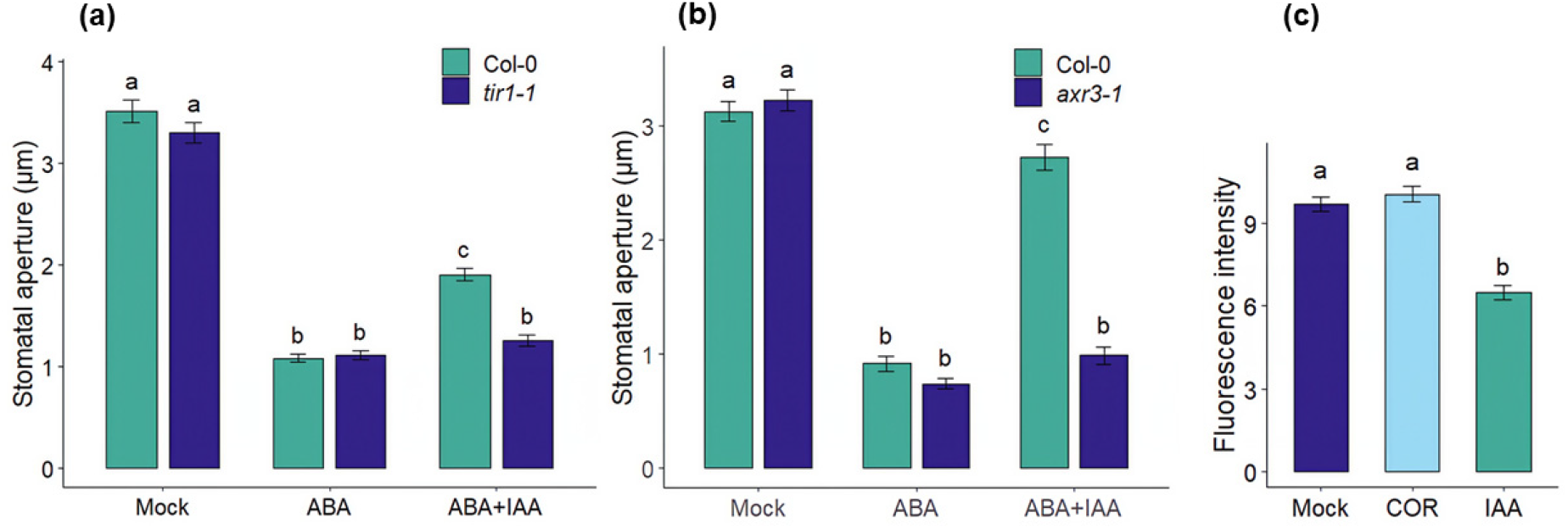
Inhibition of stomatal closure by IAA requires the presence of a TIR1/AFB–AUX/IAA signalling module. IAA (10 μM) could not inhibit stomatal closure induced by ABA (20 μM) in (a) *tir1-1* (b) *axr3-1* mutants. (c) IAA (10 μM) but not COR (1.56 μM) decreased DII-VENUS fluorescence in guard cells. Stomatal apertures were measured 1 h after ABA treatment. Different letters indicate significant differences at p < 0.05 in (a and b) (two-way ANOVA) and (c) (one-way ANOVA, Tukey′s test). Error bars represent SE from three (a and b) or two (c) independent trials, n=40 per trial in (a) and (b) and at least 40 in (c).

Next, we tested if IAA can inhibit ABA-induced ROS production in guard cells, since auxin-induced stomatal opening in the dark was shown to involve a decrease of ROS in guard cells of *Vicia faba* (Song et al., 2006), and coronatine inhibition of ABA-induced stomatal closure in Arabidopsis was associated with decreased ROS synthesis (Toum et al., 2016). We found that IAA inhibited induction of ROS by ABA (Fig. 3a), and that *rbohd*, a mutant lacking NADPH oxidase D, involved ABA-induced ROS synthesis in guard cells (Kwak, 2003), does not inhibit ABA-induced stomatal closure in response to IAA (Fig. 3b).

**Figure 3.**
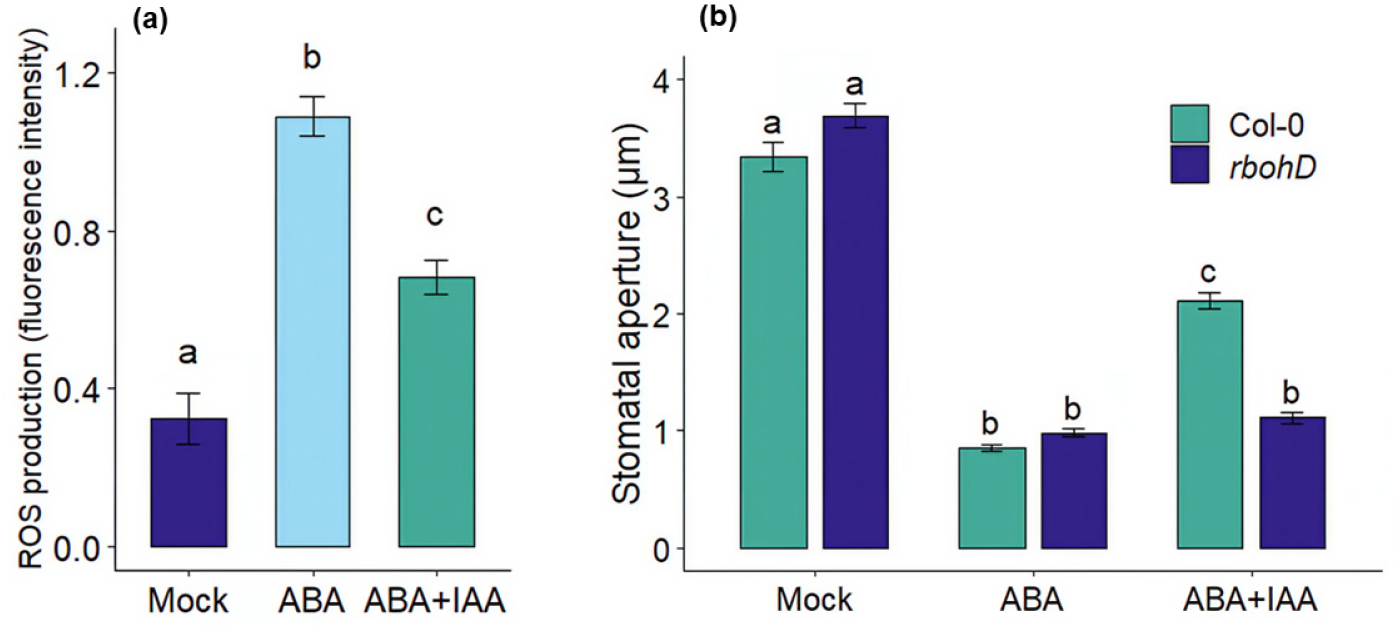
Auxin inhibits ABA-induced ROS synthesis in guard cells. (a) IAA (10 μM) inhibited ABA-induced ROS production in Col-0 guard cells. (b) IAA (10 μM) failed to inhibit stomatal closure by ABA (20 μM) in *rbohD*. In (a) epidermal peels were incubated for 20 min with the different treatments. In (b) stomatal apertures were measured 1 h after ABA treatment. Different letters indicate significant differences at p < 0.05 in (a) (one-way ANOVA, Tukey′s test) and in (b) (two-way ANOVA, Tukey′s test). Error bars represent SE from three independent trials, n=40 per trial.

Given that auxin can activate AHA2 by phosphorylation (Li et al., 2021), and that AHA2 has been linked to stomatal responses (Wong et al., 2020; Pei et al., 2022), we analysed auxin stomatal response of *aha2-5*, an AHA2 mutant allele reported as insensitive to high-temperature-induced stomatal opening (Kostaki et al., 2020). We observed that in *aha2-5* IAA cannot inhibit ABA-induced stomatal closure at 22°C. However, inhibition was restored at 35°C, indicating a possible temperature dependent role of AHA2 in mediating IAA signalling in guard cells (Fig 4a).

**Figure 4.**
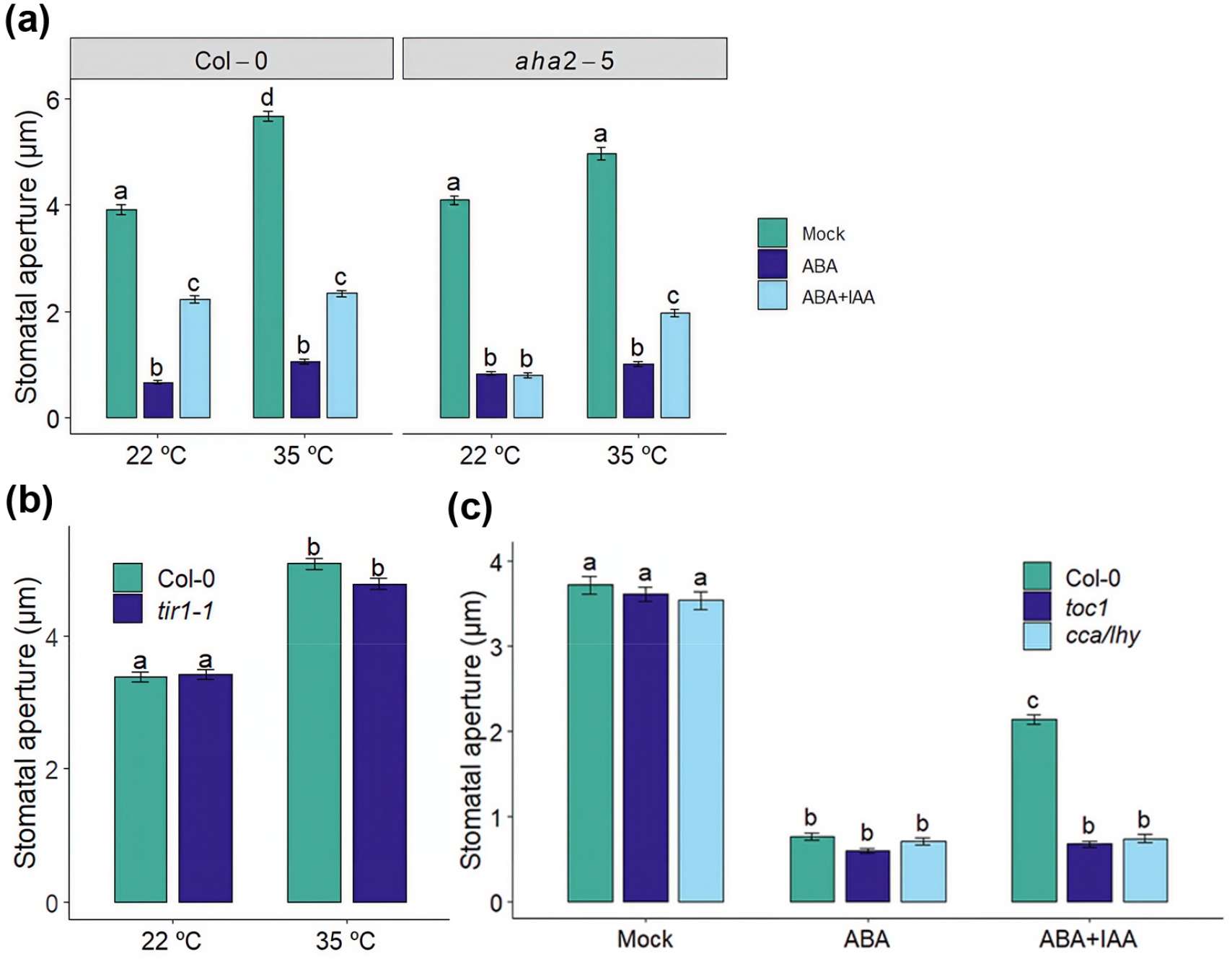
Auxin inhibition of ABA-induced stomata closure displays a temperature-dependent requirement of AHA2. (a) IAA (10 μM) did not inhibit stomatal closure induced by ABA (20 μM) in *aha2-5* mutant at 22°C, but it did at 35°C. (b) High-temperature-induced stomatal opening was not affected in *tir1-1* mutant. (c) IAA (10 μM) did not inhibit stomatal closure induced by ABA (20 μM) in circadian clock mutants *toc1* and *cca1/lhy*. Stomatal apertures were measured 1 h after ABA treatment. Different letters indicate significant differences at p < 0.05 in (a) and (b) (two-way ANOVA, Tukey′s test). In (a) data was analysed using a generalised linear mixed model (GLMM). The model included genotype, treatment, and temperature, as well as their interactions, as fixed effects. Error bars represent SE from three independent trials, n=40 per trial (a,b,c).

To find out if auxin might mediate stomatal response to high temperature, we tested *tir1-1* and found a similar stomatal aperture at 35°C than in Col-0, suggesting that IAA is not involved in this response in spite of mediating long-term adaptation to high temperature, as previously discussed (Gommers, 2020). Circadian clock responses involve the regulation of H^+^-ATPase activity (Kamrani et al., 2022), and the central regulator TOC1 has been linked to stomatal responses to drought (Legnaioli et al., 2009), while double *cca1/lhy* mutant also affected in clock regulators displays enhanced susceptibility to *P. syringae* pv. *tomato* by leaf spray but not by infiltration, suggesting that it is insensitive to coronatine in stomata (Zhang et al., 2013). Therefore, we assayed the stomatal response of *toc1* and *cca1/lhy* mutants and found that both of them are not affected by IAA in ABA-induced stomatal closure (Fig. 4c).

Auxin transport into guard cells appears to be required for stomatal sensitivity to IAA (Cho et al., 2012), and IAA concentration inside Arabidopsis guard cell protoplasts is quickly reduced in response to ABA (Jin et al., 2013), suggesting that control of its concentration inside guard cells plays a relevant physiological role. Therefore, we tested if mutants affected in auxin transport or synthesis are affected in the capacity of IAA to reduce ABA-induced stomatal closure. We found that IAA transport mutants *aux1-7* (Fig. 5a) and *pils5-2/2-5*, as well as PILS5 overexpressor (PILS5 OE) displayed reduced stomatal sensitivity to IAA (Fig. 5b). A similar insensitivity was observed in the triple *yuc5/8/9* mutant affected in auxin biosynthesis, as well as in YUCCA8 overexpressor (YUC8 OE) (Fig. 5c). Since *phyB-9* mutants display increased auxin accumulation and sensitivity (Casal & Qüesta, 2018; Legris et al., 2019), we analysed if their stomatal response to ABA is affected by IAA and found that it was not (Fig. 5d). These results provide evidence that stomatal sensitivity to auxin is tightly regulated by its own concentration or subcellular localisation.

**Figure 5.**
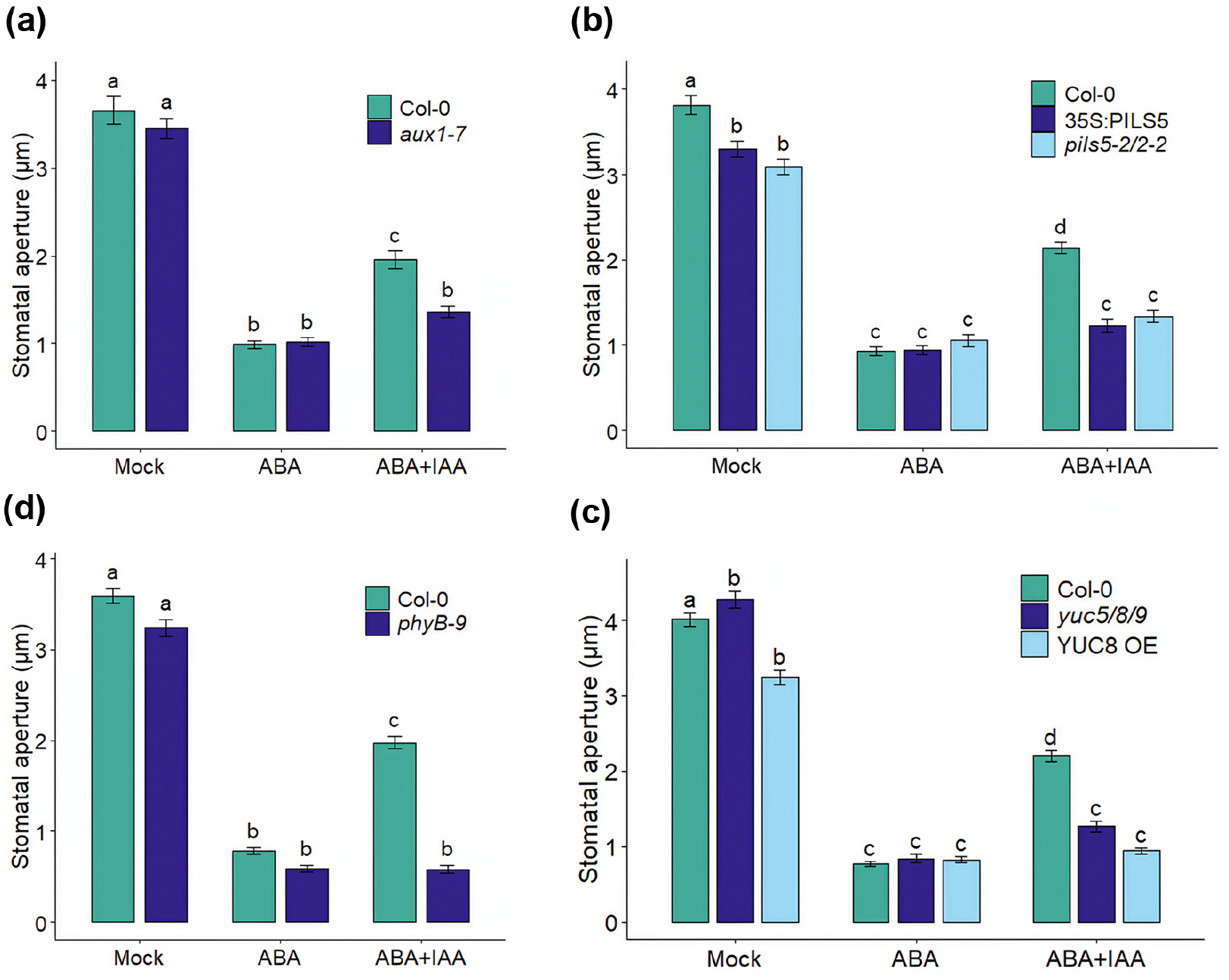
The ability of IAA to inhibit stomatal closure is reduced in mutants affected in auxin synthesis, transport and sensitivity. IAA (10 μM) showed reduced capacity to prevent stomatal closure by ABA (20 μM) in *aux1-7* (a), double *pils 5-2/2-2* mutant and 35S:PILS5 (b), triple *yuc5/yuc8/yuc9* mutant and YUCCA8 OE (YUC8 OE) (c) and *phyB-9* (d). Stomatal apertures were measured 1 h after ABA treatment. Different letters indicate significant differences at p < 0.05 (two-way ANOVA, Tukey′s test). Error bars represent SE from three independent trials, n=40 per trial (a, b, c, d).

Although the antagonistic role of ABA and IAA on stomatal aperture suggests that auxin might enhance growth at the expense or water use efficiency (WUE), numerous reports have found that auxin is positively associated with drought stress tolerance (Sharma et al., 2023). Since we have performed all previous experiments under well-watered conditions, we tested if stomatal sensitivity to auxin was affected in response to mild drought, and found that after a week of drought treatment the ability of IAA to diminish ABA-induced stomatal closure was reduced (Fig. 6a). Since there is clear evidence that ABA treatment, and therefore possibly also drought, triggers extensive chromatin remodelling and transcriptional changes in Arabidopsis guard cells (Pandey et al., 2010; Seller & Schroeder, 2023), we hypothesised that the observed variation in guard cell auxin sensitivity in response to drought might be due to underlying transcriptional changes. Consistently with this possibility, several mutants affected in the gene silencing machinery (*ago4-1, rdr2, rdr6, dcl2/3/4* and DNA demethylase triple mutant *rdd* (*ros1/dml2/dml3*), were all unresponsive to IAA in ABA-induced stomatal closure (Fig. 6b). These results indicate that the stomatal response of Arabidopsis to auxin is affected by mild drought, likely through ABA-induced transcriptional changes.

**Figure 6.**
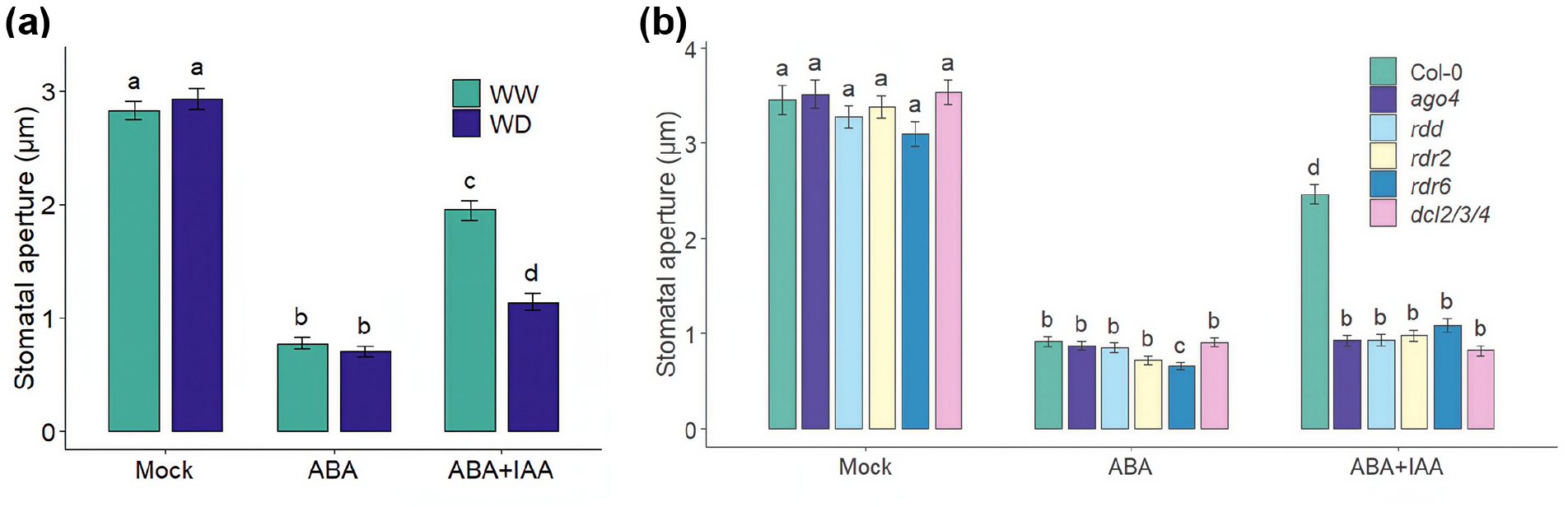
Water deficit modulates auxin sensitivity in guard cells. (a) IAA (10 μM) caused decreased inhibition ABA-induced stomatal closure (20 μM) when a mild water deficit (WD) was applied relative to well watered controls (WW) in Col-0. For water deficit treatment, plants were subjected to mild drought for one week. (b) *ago4-1, rdr2, rdr6, dcl2/3/4* and triple DNA demethylase mutant *rdd*, all affected in gene silencing, were not affected by IAA (10 μM) in promotion of stomatal closure by ABA(20 μM). Different letters indicate significant differences at p < 0.05 (two-way ANOVA, Tukey′s test). Error bars represent SE from three independent trials, n=40 per trial.

Lastly, we investigated if there is natural variation in stomatal sensitivity to auxin. For this purpose, we tested the capacity of IAA to antagonise ABA-induced stomatal closure in Col-0, Wei-0, Li-7, and St-0 ecotypes, and found that the last one was unresponsive to auxin (Fig. 7a). As St-0 was previously found to display a smaller reduction in total leaf area in response to mild drought (Bouchabke et al., 2008), and that the rapid closure of stomata has been linked to sustained growth under mild drought in Arabidopsis (Chen et al., 2021), we analysed the stomatal response to auxin of An-1 and Cvi-0, previously shown to be less affected by mild drought than Col-0, and of Mt-0, which displays the opposite behaviour (Bouchabke et al., 2008; Vile et al., 2012; Clauw et al., 2015). We observed that drought-tolerant ecotypes An-1 and Cvi-0 were insensitive to IAA in stomatal response, while in Mt-0 IAA sensitivity was slightly enhanced (Fig. 7b,c). These results show that stomatal sensitivity to auxin in Arabidopsis display natural variation and suggest that it might be linked to tolerance to mild drought.

**Figure 7.**
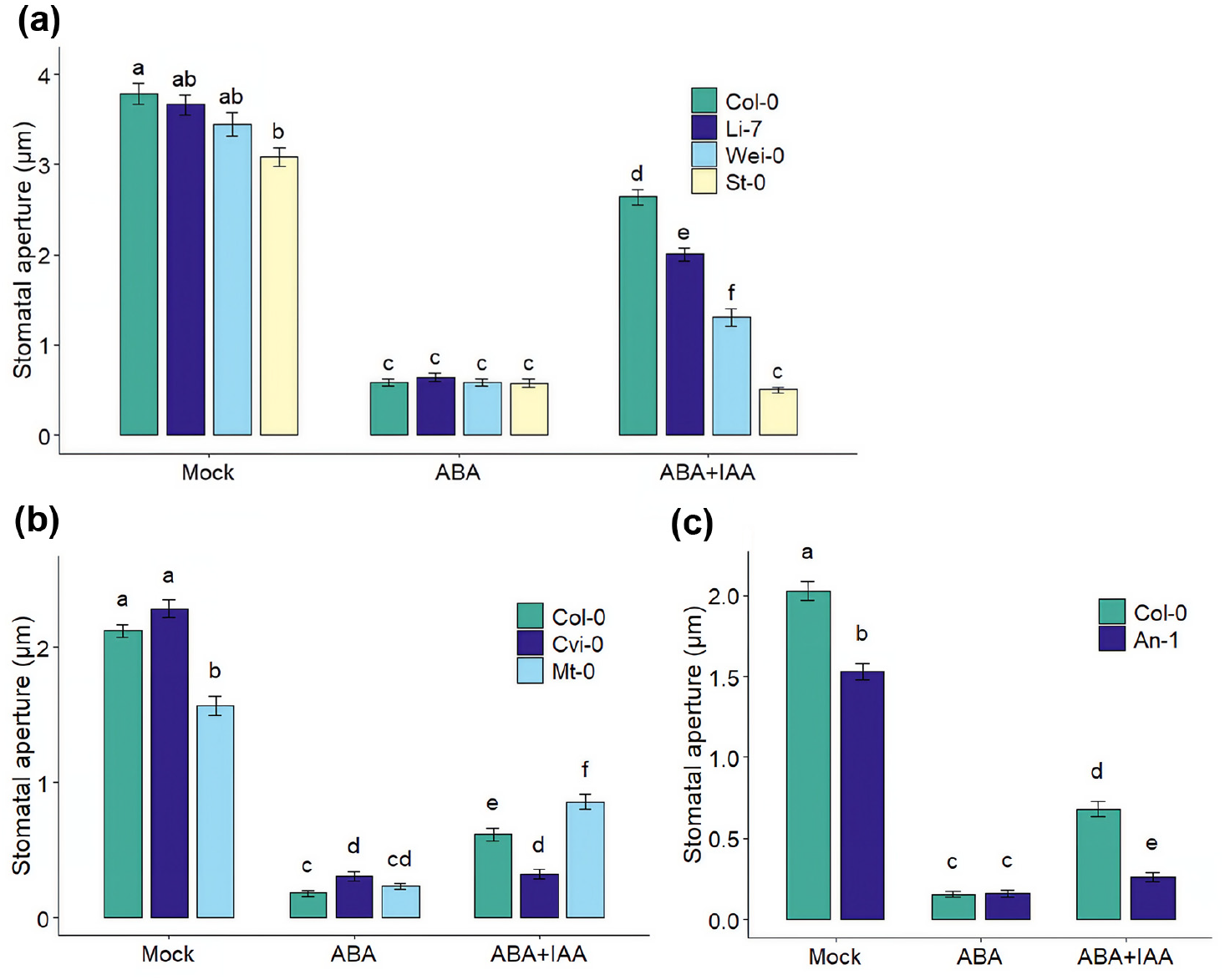
Natural variation of IAA sensitivity in guard cells. IAA (10 μM) did not inhibit ABA-induced stomatal closure (20 μM) in St-0 (a), Cvi-0 (b) and An-1 (c) ecotypes, but it did in Wei-0, Li-7 and Mt-0. Stomatal apertures were measured 1 h after ABA treatment. Different letters indicate significant differences at p < 0.05 (two-way ANOVA, Tukey′s test). Error bars represent SE from two independent trials, n=40 per trial (a, b, c).

## Discussion

Our results provide evidence that auxin inhibition of ABA-induced stomatal closure is mediated by the TIR1/AFB–AUX/IAA module. It also involves the inhibition of ROS synthesis and requires the participation of the AHA2 H^+^-ATPase. Additionally, we found that auxin sensitivity in guard cells can be affected by auxin transport, synthesis, mild drought, circadian rhythm and presents natural variation in Arabidopsis.

Although our experiments do not allow to distinguish whether TIR1/AFB–AUX/IAA involvement in stomatal closure requires or not transcriptional events, cyclic nucleotides produced upon auxin treatment by TIR1/AFB are good candidates to directly mediate its effect on stomata, perhaps independently from transcription. cAMP can promote stomatal opening in the dark (Curvetto et al., 1994), and of reversing the inhibition of guard cell inward K^+^ currents caused by ABA (Jin & Wu, 1999). cGMP similarly promotes stomatal opening, and both a guanylate cyclase inhibitor and a cGMP antagonist blocked the induction of stomatal opening in the dark by the auxin indolyl-3butyric acid (IBA) (Cousson & Vavasseur, 1998; Cousson, 2001). Therefore, both might mediate auxin effect on stomatal aperture. Although our results show an involvement of TIR1/AFB–AUX/IAA in inhibition of ABA-induced stomatal closure by IAA, the very rapid responses elicited by this hormone on electric currents in guard cells (Lohse & Hedrich, 1992; Blatt & Thiel, 1994) indicate that other fast perception mechanisms might be involved. ABP1 could also mediate auxin effect in guard cells, as its overexpression enhanced inwardly and outwardly rectifying K^+^ currents in a dose-dependent manner in response to IAA in tobacco guard cells (Bauly et al., 2000). ABP1 activates TMK kinases, which phosphorylate AHAs and promote cell-wall acidification and hypocotyl cell elongation in Arabidopsis (Lin et al., 2021). A similar regulation of H^+^-ATPase activity might also exist in guard cells, as the TMK1 promoter is active in them (Yang et al., 2021). This study also found that phosphorylation by TMK1 in Thr-321 inhibits the phosphatase activity of ABI2, involved in ABA signalling. While this report showed that *tmk1* mutants have increased stomatal aperture, a recent study found an opposite phenotype (Monzer et al., 2025), and therefore the role of TMK1 in guard cells is unclear. TIR1/AFB–AUX/IAA might also mediate auxin effect through transcriptional regulation of auxin-induced SAUR proteins, which inhibit PP2C D clade phosphatases that dephosphorylate and inactivate AHA2 in guard cells (Wong et al., 2020). Thus, auxin activation of SAURs should relieve the inhibitory effect of PP2C D phosphatases on AHA2 and promote stomatal aperture. While in this study a direct regulation by auxin was not demonstrated, the role of SAURs in mediating hypocotyl elongation by IAA is well established (Fendrych et al., 2016). As has been proposed for regulation of growth by auxin in roots and hypocotyls (Vanneste et al., 2025), in stomata IAA-regulated H^+^-ATPase activity might also be controlled by transcriptional and non-transcriptional regulation in different time frames.

The observed inhibitory effect of auxin on ABA-induced ROS synthesis in guard cells is consistent with their important role as second messengers of this hormone in stomata (Cho et al., 2009). ABA-induced ROS are synthesised by NADPH oxidases, and lead to the activation of the plasma membrane Ca^2+^-permeable channels and SLAC1 anion channel (Waadt et al., 2022), as well as the inhibition of plasma membrane H^+^-ATPase activity (Zhang et al., 2004). In agreement with these results, the Ca^2+^ buffer BAPTA antagonises auxin effect in stomata (Cousson & Vavasseur, 1998). Previously, it was shown that the bacterial toxin coronatine, which facilitates *P. syringae* pv. *tomato* penetration through stomata, also inhibits ABA-induced ROS production (Toum et al., 2016), which suggests that auxin and this toxin might act through shared signalling targets. Interestingly, certain bacterial effectors also inhibit flagellin-triggered ROS synthesis in guard cells (Melotto et al., 2017). In addition, Arabidopsis RPM1-INTERACTING4 (RIN4), a negative regulator of plant innate immunity and target of bacterial effectors, is required for coronatine effect on stomatal closure and can interact with AHA1 and AHA2, suggesting that coronatine regulates H^+^-ATPase activity (Liu et al., 2009). When RIN4 is activated by phosphorylation, it promotes H^+^-ATPase activity and inhibits flagellin-triggered ROS synthesis in Arabidopsis (Lee et al., 2015). The fact that different pathogens appear to target ROS synthesis in guard cells to inhibit stomatal closure, indicates the crucial role of ROS as second messengers for the regulation of stomatal aperture, through the control of ion channels and AHA activity.

The stomatal insensitivity of *aha2-5* to IAA is consistent with a role of AHA2 in mediating auxin effect on guard cells, at least at 22°C. The fact that *aha2-5* responds to auxin at 35°C might imply that at high temperature AHAs are regulated differently, so that different members of the family could mediate the effect of auxin depending on the temperature. Our results show that IAA can inhibit stomatal closure induced by ABA similarly at 22°C and 35°C, but the fact that *tir1-1* show a similar response to high temperature as Col-0 suggests that, as previously proposed (Gommers, 2020), auxin does not mediate high-temperature-induced stomatal opening.

The circadian clock plays in important role in the regulation of stomatal aperture (Casal & Qüesta, 2018), as well as in the optimisation of water use (Simon et al., 2020; Kamrani et al., 2022), a process that might involve the regulation of guard cell H^+^-ATPases. Our finding that *toc1* and *cca1/lhy* do not respond to auxin might imply that these mutants have less active AHAs. Consistent with the possibility that the circadian clock regulate H^+^-ATPase activity in guard cells, guard-cell-specific overexpression of CCA1 in Arabidopsis led to increased biomass accumulation under fully watered and mild drought conditions (Hassidim et al., 2017), an effect also achieved by guard-cell-specific overexpression of AHA2 (Wang et al., 2014). Our observation that *phyB-9* is insensitive in stomata to auxin also supports the possibility of a link between sensitivity to this hormone and water use. PHYB was shown to have a negative impact on water use, as *phyB-5* mutant displays increased long term WUE as assessed by carbon isotope discrimination (Boccalandro et al., 2009). The reduced ability of auxin to inhibit stomatal closure in ecotypes St-0, An-1 and Cvi-0, previously shown to be less affected in their growth than Col-0 by mild drought, but not in Mt-0, which displays an opposite response to mild drought (Bouchabke et al., 2008; Vile et al., 2012; Clauw et al., 2015), also lends support to the existence of a possible link between stomatal sensitivity to auxin and water use. Col-0 stomatal sensitivity to auxin was reduced by mild drought treatment, which might reflect possible transcriptional changes in guard cell elicited by water stress (Virlouvet & Fromm, 2015; Auler et al., 2021; Seller & Schroeder, 2023). The latter study reported that *AHA2* and *AHA5* transcripts were downregulated in guard cells by ABA treatment (*AHA1* was not studied), which might contribute to explain decreased stomatal sensitivity to IAA if such transcriptional changes were sustained in time during water stress, possibly through modifications in chromatin accesibility like those reported in the same work.

Apart from *AHA1* and *AHA2* expression, ABA also reduces IAA concentration in guard cells of Arabidopsis and *Brassica napus* (Jin et al., 2013; Zhu & Assmann, 2017). Thus, auxin metabolism and dynamics within these cells possibly play a relevant role, and consistently, we observed that affecting its synthesis or transport by mutation or transgenesis reduced its stomatal sensitivity. Auxin stomatal sensitivity seems therefore to be tightly regulated, as mutation affecting the gene silencing machinery also reduces it, although this could be due to either direct or indirect effects on guard cells (*e*.*g*. by affecting auxin transport in leaves).

Our results do not allow us to determine if auxin does play a regulatory role on stomata *in planta, i*.*e*. if actual auxin changes within or in the vicinity of guard cells can modulate stomatal aperture. It might also be possible that changes in auxin stomatal sensitivity are a consequence and not a cause of, for example, drought-induced changes in the expression of guard cell ion channels or AHAs. Nevertheless, it is possible to speculate that auxin acts as a signal that antagonises stomatal closure when conditions are favourable for carbon fixation. For example, auxin sensitivity was shown to be enhanced by glucose availability (Mishra et al., 2022), and also the circadian clock modulates multiple auxin responses (Covington & Harmer, 2007), which are in part regulated by direct interaction of CCA1/LHY with Aux/IAA proteins (Xue et al., 2020). Thus, changing sensitivity to auxin might contribute to changes in stomatal aperture during the day. Natural variation of growth sensitivity in response to mild drought in Arabidopsis has been associated with the speed of stomatal response (Chen et al., 2021), which our results suggest that might be associated to stomatal auxin sensitivity. While further studies are needed to confirm this hypothesis, the existence of natural variation in stomal response to auxin suggests that this trait has been subjected to selective pressures and might provide a new target for the selection of drought tolerant plants.

## Acknowledgements

This work was supported by grants PICT 2014-3286, PICT-2017-2075, 11220200102958CO and PIP 00903-2016. G.E.G, L.T, and F.L.P Career Investigators of CONICET, C.A.R was supported by a doctoral scholarship from CONICET. We thank technical assistance from María Victoria Repetto with microscopy and Lucas Kreiman for help with statistical analysis.

## Author contributions

G.E.G., L.T., and F.L.P. conceived the project. G.E.G., L.T., C.A.R. and L.P. performed the experiments. C.A.R. performed statistical analysis. G.E.G. wrote the manuscript and L.T., C.A.R. and L.P. revised it. All authors approved the final version.

## References

Abuzeineh A, Abualia R. 2023. Unlocking Auxin’s Potential: How Plants Use Auxin Transport to Beat Drought. Biomedical Journal of Scientific & Technical Research 51: 42712–42715.

Alabadí D, Yanovsky MJ, Más P, Harmer SL, Kay SA. 2002. Critical Role for CCA1 and LHY in Maintaining Circadian Rhythmicity in Arabidopsis. Current Biology 12: 757–761.

Aliniaeifard S, Van Meeteren U. 2014. Natural variation in stomatal response to closing stimuli among Arabidopsis thaliana accessions after exposure to low VPD as a tool to recognize the mechanism of disturbed stomatal functioning. Journal of Experimental Botany 65: 6529–6542.

Alonso-Blanco C, Andrade J, Becker C, Bemm F, Bergelson J, Borgwardt KMM, Cao J, Chae E, Dezwaan TMM, Ding W, et al. 2016. 1,135 Genomes Reveal the Global Pattern of Polymorphism in Arabidopsis thaliana. Cell 166: 481–491.

Ang ACH, Østergaard L. 2023. Save your TIRs - more to auxin than meets the eye. The New Phytologist 238: 971–976.

Auler PA, Nogueira do Amaral M, Bolacel Braga EJ, Maserti B. 2021. Drought stress memory in rice guard cells: Proteome changes and genomic stability of DNA. Plant physiology and biochemistry: PPB 169: 49–62.

Barbez E, Kubeš M, Rolčík J, Béziat C, Pěnčík A, Wang B, Rosquete MR, Zhu J, Dobrev PI, Lee Y, et al. 2012. A novel putative auxin carrier family regulates intracellular auxin homeostasis in plants. Nature 485: 119–122.

Bauly JM, Sealy IM, Macdonald H, Brearley J, Dröge S, Hillmer S, Robinson DG, Venis MA, Blatt MR, Lazarus CM, et al. 2000. Overexpression of Auxin-Binding Protein Enhances the Sensitivity of Guard Cells to Auxin1. Plant Physiology 124: 1229–1238.

Blakeslee JJ, Spatola Rossi T, Kriechbaumer V. 2019. Auxin biosynthesis: spatial regulation and adaptation to stress (C Raines, Ed.). Journal of Experimental Botany 70: 5041–5049.

Blatt MR, Thiel G. 1994. K+ channels of stomatal guard cells: bimodal control of the K+ inward-rectifier evoked by auxin. The Plant Journal 5: 55–68.

Boccalandro HE, Rugnone ML, Moreno JE, Ploschuk EL, Serna L, Yanovsky MJ, Casal JJ. 2009. Phytochrome B Enhances Photosynthesis at the Expense of Water-Use Efficiency in Arabidopsis. Plant Physiology 150: 1083–1092.

Bouchabke O, Chang F, Simon M, Voisin R, Pelletier G, Durand-Tardif M. 2008. Natural Variation in Arabidopsis thaliana as a Tool for Highlighting Differential Drought Responses (J Kroymann, Ed.). PLoS ONE 3: e1705.

Brooks M E, Kristensen K, Benthem K J, van, Magnusson A, Berg C W, Nielsen A, Skaug H J, Mächler M, Bolker B M. 2017. glmmTMB Balances Speed and Flexibility Among Packages for Zero-inflated Generalized Linear Mixed Modeling. The R Journal 9: 378.

Brunoud G, Wells DM, Oliva M, Larrieu A, Mirabet V, Burrow AH, Beeckman T, Kepinski S, Traas J, Bennett MJ, et al. 2012. A novel sensor to map auxin response and distribution at high spatio-temporal resolution. Nature 482: 103–106.

Casal JJ, Qüesta JI. 2018. Light and temperature cues: multitasking receptors and transcriptional integrators. The New Phytologist 217: 1029–1034.

Chen Q, Dai X, De-Paoli H, Cheng Y, Takebayashi Y, Kasahara H, Kamiya Y, Zhao Y. 2014. Auxin Overproduction in Shoots Cannot Rescue Auxin Deficiencies in Arabidopsis Roots. Plant and Cell Physiology 55: 1072–1079.

Chen Y, Dubois M, Vermeersch M, Inzé D, Vanhaeren H. 2021. Distinct cellular strategies determine sensitivity to mild drought of Arabidopsis natural accessions. Plant Physiology 186: 1171–1185.

Chen H, Qi L, Zou M, Lu M, Kwiatkowski M, Pei Y, Jaworski K, Friml J. 2025. TIR1-produced cAMP as a second messenger in transcriptional auxin signalling. Nature 640: 1011–1016.

Cho D, Shin D, Jeon BW, Kwak JM. 2009. ROS-Mediated ABA Signaling. Journal of Plant Biology 52: 102–113.

Cho D, Villiers F, Kroniewicz L, Lee S, Seo YJ, Hirschi KD, Leonhardt N, Kwak JM. 2012. Vacuolar CAX1 and CAX3 Influence Auxin Transport in Guard Cells via Regulation of Apoplastic pH. Plant Physiology 160: 1293–1302.

Clauw P, Coppens F, De Beuf K, Dhondt S, Van Daele T, Maleux K, Storme V, Clement L, Gonzalez N, Inzé D. 2015. Leaf Responses to Mild Drought Stress in Natural Variants of Arabidopsis. Plant Physiology 167: 800–816.

Cousson A. 2001. Pharmacological evidence for the implication of both cyclic GMP-dependent and - independent transduction pathways within auxin-induced stomatal opening in Commelina communis (L.). Plant Science 161: 249–258.

Cousson A, Vavasseur A. 1998. Putative involvement of cytosolic Ca2+ and GTP-binding proteins in cyclic-GMP-mediated induction of stomatal opening by auxin in Commelina communis L. Planta 206: 308–314.

Covington MF, Harmer SL. 2007. The Circadian Clock Regulates Auxin Signaling and Responses in Arabidopsis. PLoS Biology 5: e222.

Curvetto N, Darjania. L, Delmastro S. 1994. Effect of 2 cAMP analogs on stomatal opening in Vicia Faba. Possible relationship with cytosolic calcium-concentration. Plant Physiology and Biochemistry 32: 365–372.

Ding Y, Fromm M, Avramova Z. 2012. Multiple exposures to drought ‘train’ transcriptional responses in Arabidopsis. Nature Communications 3: 740.

Dunleavy PJ, Ladley PD. 1995. Stomatal responses of Vicia faba L. to indole acetic acid and abscisic acid. Journal of Experimental Botany 46: 95–100.

Fendrych M, Leung J, Friml J. 2016. TIR1/AFB-Aux/IAA auxin perception mediates rapid cell wall acidification and growth of Arabidopsis hypocotyls. eLife 5: e19048.

Fuglsang AT, Guo Y, Cuin TA, Qiu Q, Song C, Kristiansen KA, Bych K, Schulz A, Shabala S, Schumaker KS, et al. 2007. Arabidopsis Protein Kinase PKS5 Inhibits the Plasma Membrane H+-ATPase by Preventing Interaction with 14-3-3 Protein. The Plant Cell 19: 1617–1634.

Gommers C. 2020. Keep Cool and Open Up: Temperature-Induced Stomatal Opening. Plant Physiology 182: 1188–1189.

Gudesblat GE, Torres PS, Vojnov AA. 2009. Xanthomonas campestris Overcomes Arabidopsis Stomatal Innate Immunity through a DSF Cell-to-Cell Signal-Regulated Virulence Factor. Plant Physiology 149: 1017– 1027.

Hartig F. 2024. DHARMa: Residual Diagnostics for Hierarchical (Multi-Level / Mixed) Regression Models. https://github.com/florianhartig/dharma

Haruta M, Burch HL, Nelson RB, Barrett-Wilt G, Kline KG, Mohsin SB, Young JC, Otegui MS, Sussman MR. 2010. Molecular characterization of mutant Arabidopsis plants with reduced plasma membrane proton pump activity. The Journal of Biological Chemistry 285: 17918–17929.

Hassidim M, Dakhiya Y, Turjeman A, Hussien D, Shor E, Anidjar A, Goldberg K, Green RM. 2017. CIRCADIAN CLOCK ASSOCIATED1 (CCA1) and the Circadian Control of Stomatal Aperture. Plant Physiology 175: 1864–1877.

Henderson IR, Zhang X, Lu C, Johnson L, Meyers BC, Green PJ, Jacobsen SE. 2006. Dissecting Arabidopsis thaliana DICER function in small RNA processing, gene silencing and DNA methylation patterning. Nature Genetics 38: 721–725.

Hentrich M, Böttcher C, Düchting P, Cheng Y, Zhao Y, Berkowitz O, Masle J, Medina J, Pollmann S. 2013. The jasmonic acid signaling pathway is linked to auxin homeostasis through the modulation of YUCCA 8 and YUCCA 9 gene expression. The Plant Journal 74: 626–637.

Hsu P-K, Dubeaux G, Takahashi Y, Schroeder JI. 2021. Signaling mechanisms in abscisic acid-mediated stomatal closure. The Plant Journal 105: 307–321.

Irving HR, Gehring CA, Parish RW. 1992. Changes in cytosolic pH and calcium of guard cells precede stomatal movements. Proceedings of the National Academy of Sciences 89: 1790–1794.

Jin X, Wang R-S, Zhu M, Jeon BW, Albert R, Chen S, Assmann SM. 2013. Abscisic acid-responsive guard cell metabolomes of Arabidopsis wild-type and gpa1 G-protein mutants. The Plant Cell 25: 4789–4811.

Jin X-C, Wu W-H. 1999. Involvement of Cyclic AMP in ABA- and Ca2−-Mediated Signal Transduction of Stomatal Regulation in Vicia faba. Plant and Cell Physiology 40: 1127–1133.

Kamrani Y, Shomali A, Aliniaeifard S, Lastochkina O, Moosavi-Nezhad M, Hajinajaf N, Talar U. 2022. Regulatory Role of Circadian Clocks on ABA Production and Signaling, Stomatal Responses, and Water-Use Efficiency under Water-Deficit Conditions. Cells 11: 1154.

Kollist H, Nuhkat M, Roelfsema MRG. 2014. Closing gaps: linking elements that control stomatal movement. The New Phytologist 203: 44–62.

Kostaki KI, Coupel-Ledru A, Bonnell VC, Gustavsson M, Sun P, McLaughlin FJ, Fraser DP, McLachlan DH, Hetherington AM, Dodd AN, et al. 2020. Guard cells integrate light and temperature signals to control stomatal aperture. Plant Physiology 182: 1404–1419.

Kwak JM. 2003. NADPH oxidase AtrbohD and AtrbohF genes function in ROS-dependent ABA signaling in Arabidopsis. The EMBO Journal 22: 2623–2633.

Le T-N, Schumann U, Smith NA, Tiwari S, Au PCK, Zhu Q-H, Taylor JM, Kazan K, Llewellyn DJ, Zhang R, et al. 2014. DNA demethylases target promoter transposable elements to positively regulate stress responsive genes in Arabidopsis. Genome Biology 15: 458.

Lee D, Bourdais G, Yu G, Robatzek S, Coaker G. 2015. Phosphorylation of the Plant Immune Regulator RPM1-INTERACTING PROTEIN4 Enhances Plant Plasma Membrane H+-ATPase Activity and Inhibits Flagellin-Triggered Immune Responses in Arabidopsis. The Plant Cell 27: 2042–2056.

Leftley N, Banda J, Pandey B, Bennett M, Voß U. 2021. Uncovering How Auxin Optimizes Root Systems Architecture in Response to Environmental Stresses. Cold Spring Harbor Perspectives in Biology 13: a040014.

Legnaioli T, Cuevas J, Mas P. 2009. TOC1 functions as a molecular switch connecting the circadian clock with plant responses to drought. The EMBO Journal 28: 3745–3757.

Legris M, Ince YÇ, Fankhauser C. 2019. Molecular mechanisms underlying phytochrome-controlled morphogenesis in plants. Nature Communications 10: 5219.

Levitt LK, Stein DB, Rubinstein B. 1987. Promotion of Stomatal Opening by Indoleacetic Acid and Ethrel in Epidermal Strips of Vicia faba L. Plant Physiology 85: 318–321.

Leyser O. 2018. Auxin Signaling. Plant Physiology 176: 465–479.

Leyser HMO, Pickett FB, Dharmasiri S, Estelle M. 1996. Mutations in the AXR3 gene of Arabidopsis result in altered auxin response including ectopic expression from the SAUR-AC1 promoter. The Plant Journal 10: 403–413.

Li L, Verstraeten I, Roosjen M, Takahashi K, Rodriguez L, Merrin J, Chen J, Shabala L, Smet W, Ren H, et al. 2021. Cell surface and intracellular auxin signalling for H+ fluxes in root growth. Nature 599: 273– 277.

Li Y, Zeng H, Xu F, Yan F, Xu W. 2022. H+-ATPases in Plant Growth and Stress Responses. Annual Review of Plant Biology 73: 495–521.

Lin W, Zhou X, Tang W, Takahashi K, Pan X, Dai J, Ren H, Zhu X, Pan S, Zheng H, et al. 2021. TMK-based cell-surface auxin signalling activates cell-wall acidification. Nature 599: 278–282.

Liu J, Elmore JM, Fuglsang AT, Palmgren MG, Staskawicz BJ, Coaker G. 2009. RIN4 Functions with Plasma Membrane H+-ATPases to Regulate Stomatal Apertures during Pathogen Attack. PLOS Biology 7: e1000139.

Lohse G, Hedrich R. 1992. Characterization of the plasma-membrane H+-ATPase from Vicia faba guard cells: Modulation by extracellular factors and seasonal changes. Planta 188: 206–214.

Marten I, Lohse G, Hedrich R. 1991. Plant growth hormones control voltage-dependent activity of anion channels in plasma membrane of guard cells. Nature 353: 758–762.

Melotto M, Underwood W, Koczan J, Nomura K, He SY. 2006. Plant Stomata Function in Innate Immunity against Bacterial Invasion. Cell 126: 969–980.

Melotto M, Zhang L, Oblessuc PR, He SY. 2017. Stomatal Defense a Decade Later. Plant Physiology 174: 561–571.

Mishra BS, Sharma M, Laxmi A. 2022. Role of sugar and auxin crosstalk in plant growth and development. Physiologia Plantarum 174: e13546.

Monroe JG, Powell T, Price N, Mullen JL, Howard A, Evans K, Lovell JT, McKay JK. 2018. Drought adaptation in Arabidopsis thaliana by extensive genetic loss-of-function. eLife 7: e41038.

Monzer A, Mazur E, Rodriguez L, Gallei M, Zou M, Smejkal M, Červeňová E, Friml J. 2025. TMK interacting network of receptor like kinases for auxin canalization and beyond. bioRxiv: 2025.02.28.640727.

Murata Y, Pei Z-M, Mori IC, Schroeder J. 2001. Abscisic Acid Activation of Plasma Membrane Ca2+ Channels in Guard Cells Requires Cytosolic NAD(P)H and Is Differentially Disrupted Upstream and Downstream of Reactive Oxygen Species Production in abi1-1 and abi2-1 Protein Phosphatase 2C Mutants. The Plant Cell 13: 2513–2523.

Pandey S, Wang R-S, Wilson L, Li S, Zhao Z, Gookin TE, Assmann SM, Albert R. 2010. Boolean modeling of transcriptome data reveals novel modes of heterotrimeric G-protein action. Molecular systems biology 6: 372.

Pei D, Hua D, Deng J, Wang Z, Song C, Wang Y, Wang Y, Qi J, Kollist H, Yang S, et al. 2022. Phosphorylation of the plasma membrane H+-ATPase AHA2 by BAK1 is required for ABA-induced stomatal closure in Arabidopsis. The Plant Cell 34: 2708–2729.

Peragine A, Yoshikawa M, Wu G, Albrecht HL, Poethig RS. 2004. SGS3 and SGS2/SDE1/RDR6 are required for juvenile development and the production of trans -acting siRNAs in Arabidopsis. Genes & Development 18: 2368–2379.

Pickett FB, Wilson AK, Estelle M. 1990. The aux1 Mutation of Arabidopsis Confers Both Auxin and Ethylene Resistance. Plant Physiology 94: 1462–1466.

Pospíšilová J. 2003. Participation of Phytohormones in the Stomatal Regulation of Gas Exchange During Water Stress. Biologia plantarum 46: 491–506.

Qi L, Kwiatkowski M, Kulich I, Chen H, Gao Y, Yun P, Li L, Shabala S, Farmer EE, Jaworski K. 2023. Guanylate cyclase activity of TIR1/AFB auxin receptors in rapid auxin responses. bioRxiv: 2023.11. 18.567481.

Reed JW, Nagpal P, Poole DS, Furuya M, Chory J. 1993. Mutations in the gene for the red/far-red light receptor phytochrome B alter cell elongation and physiological responses throughout Arabidopsis development. The Plant Cell 5: 147–157.

Ruegger M, Dewey E, Gray WM, Hobbie L, Turner J, Estelle M. 1998. The TIR1 protein of Arabidopsis functions in auxin response and is related to human SKP2 and yeast grr1p. Genes & Development 12: 198– 207.

Salehin M, Li B, Tang M, Katz E, Song L, Ecker JR, Kliebenstein DJ, Estelle M. 2019. Auxin-sensitive Aux/IAA proteins mediate drought tolerance in Arabidopsis by regulating glucosinolate levels. Nature Communications 10: 4021.

Seller CA, Schroeder JI. 2023. Distinct guard cell–specific remodeling of chromatin accessibility during abscisic acid– and CO2-dependent stomatal regulation. Proceedings of the National Academy of Sciences 120: e2310670120.

Shani E, Salehin M, Zhang Y, Sanchez SE, Doherty C, Wang R, Mangado CC, Song L, Tal I, Pisanty O, et al. 2017. Plant Stress Tolerance Requires Auxin-Sensitive Aux/IAA Transcriptional Repressors. Current Biology 27: 437–444.

Sharma A, Gupta A, Ramakrishnan M, Ha CV, Zheng B, Bhardwaj M, Tran L-SP. 2023. Roles of abscisic acid and auxin in plants during drought: A molecular point of view. Plant Physiology and Biochemistry 204: 108129.

Simon NML, Graham CA, Comben NE, Hetherington AM, Dodd AN. 2020. The Circadian Clock Influences the Long-Term Water Use Efficiency of Arabidopsis1 [OPEN]. Plant Physiology 183: 317–330.

Snaith PJ, Mansfield TA. 1982. Control of the CO2 Responses of Stomata by Indol-3ylacetic Acid and Abscisic Acid. Journal of Experimental Botany 33: 360–365.

Song X-G, She X-P, He J-M, Huang C, Song T. 2006. Cytokinin- and auxin-induced stomatal opening involves a decrease in levels of hydrogen peroxide in guard cells of Vicia faba. Functional Plant Biology 33: 573.

Strayer C, Oyama T, Schultz TF, Raman R, Somers DE, Más P, Panda S, Kreps JA, Kay SA. 2000. Cloning of the Arabidopsis Clock Gene TOC1, an Autoregulatory Response Regulator Homolog. Science 289: 768–771.

Tanaka Y. 2006. Cytokinin and auxin inhibit abscisic acid-induced stomatal closure by enhancing ethylene production in Arabidopsis. Journal of Experimental Botany 57: 2259–2266.

Torres MA, Dangl JL, Jones JDG. 2002. Arabidopsis gp91phox homologues AtrbohD and AtrbohF are required for accumulation of reactive oxygen intermediates in the plant defense response. Proceedings of the National Academy of Sciences of the United States of America 99: 517–522.

Toum L, Torres PS, Gallego SM, Benavídes MP, Vojnov AA, Gudesblat GE. 2016. Coronatine Inhibits Stomatal Closure through guard-cell-specific Inhibition of NADPH Oxidase-Dependent ROS Production. Frontiers in Plant Science 7: 1–12.

Tresas T, Isaioglou I, Roussis A, Haralampidis K. 2025. A Brief Overview of the Epigenetic Regulatory Mechanisms in Plants. International Journal of Molecular Sciences 26: 4700.

Ueno K, Kinoshita T, Inoue S, Emi T, Shimazaki K. 2005. Biochemical Characterization of Plasma Membrane H+-ATPase Activation in Guard Cell Protoplasts of Arabidopsis thaliana in Response to Blue Light. Plant and Cell Physiology 46: 955–963.

Vanneste S, Pei Y, Friml J. 2025. Mechanisms of auxin action in plant growth and development. Nature Reviews. Molecular Cell Biology.

Vile D, Pervent M, Belluau M, Vasseur F, Bresson J, Muller B, Granier C, Simonneau T. 2012. Arabidopsis growth under prolonged high temperature and water deficit: independent or interactive effects? Plant, Cell & Environment 35: 702–718.

Virlouvet L, Fromm M. 2015. Physiological and transcriptional memory in guard cells during repetitive dehydration stress. New Phytologist 205: 596–607.

Waadt R, Seller CA, Hsu P-K, Takahashi Y, Munemasa S, Schroeder JI. 2022. Plant hormone regulation of abiotic stress responses. Nature Reviews. Molecular Cell Biology 23: 680–694.

Wang Y, Noguchi K, Ono N, Inoue S, Terashima I, Kinoshita T. 2014. Overexpression of plasma membrane H+ -ATPase in guard cells promotes light-induced stomatal opening and enhances plant growth. Proceedings of the National Academy of Sciences 111: 533–538.

Wickham H. 2016. Programming with ggplot2. In: Wickham H, ed. ggplot2: Elegant Graphics for Data Analysis. Cham: Springer International Publishing, 241–253.

Wong JH, Klejchová M, Snipes SA, Nagpal P, Bak G, Wang B, Dunlap S, Park MY, Kunkel EN, Trinidad B, et al. 2020. SAUR proteins and PP2C.D phosphatases regulate H+-ATPases and K+ channels to control stomatal movements. Plant Physiology 185: 256–273.

Xiao-Ping S, Xi-Gui S. 2006. Cytokinin- and auxin-induced stomatal opening is related to the change of nitric oxide levels in guard cells in broad bean. Physiologia Plantarum 128: 569–579.

Xie Z, Johansen LK, Gustafson AM, Kasschau KD, Lellis AD, Zilberman D, Jacobsen SE, Carrington JC. 2004. Genetic and Functional Diversification of Small RNA Pathways in Plants (Detlef Weigel, Ed.). PLoS Biology 2: e104.

Xu F, He S, Zhang J, Mao Z, Wang W, Li T, Hua J, Du S, Xu P, Li L, et al. 2018. Photoactivated CRY1 and phyB Interact Directly with AUX/IAA Proteins to Inhibit Auxin Signaling in Arabidopsis. Molecular Plant 11: 523–541.

Xu X, Liu H, Praat M, Pizzio GA, Jiang Z, Driever SM, Wang R, Van De Cotte B, Villers SLY, Gevaert K, et al. 2025. Stomatal opening under high temperatures is controlled by the OST1-regulated TOT3-AHA1 module. Nature Plants 11: 105–117.

Xue X, Sun K, Zhu Z. 2020. CIRCADIAN CLOCK ASSOCIATED 1 gates morning phased auxin response in Arabidopsis thaliana. Biochemical and Biophysical Research Communications 527: 935–940.

Yamauchi S, Takemiya A, Sakamoto T, Kurata T, Tsutsumi T, Kinoshita T, Shimazaki K. 2016. The Plasma Membrane H+-ATPase AHA1 Plays a Major Role in Stomatal Opening in Response to Blue Light1. Plant Physiology 171: 2731–2743.

Yang J, He H, He Y, Zheng Q, Li Q, Feng X, Wang P, Qin G, Gu Y, Wu P, et al. 2021. TMK1-based auxin signaling regulates abscisic acid responses via phosphorylating ABI1/2 in Arabidopsis. Proceedings of the National Academy of Sciences 118: e2102544118.

Yu Z, Zhang F, Friml J, Ding Z. 2022. Auxin signaling: Research advances over the past 30 years. Journal of Integrative Plant Biology 64: 371–392.

Zhang X, Wang H, Takemiya A, Song C, Kinoshita T, Shimazaki K. 2004. Inhibition of Blue Light-Dependent H+ Pumping by Abscisic Acid through Hydrogen Peroxide-Induced Dephosphorylation of the Plasma Membrane H+-ATPase in Guard Cell Protoplasts. Plant Physiology 136: 4150–4158.

Zhang C, Xie Q, Anderson RG, Ng G, Seitz NC, Peterson T, McClung CR, McDowell JM, Kong D, Kwak JM, et al. 2013. Crosstalk between the Circadian Clock and Innate Immunity in Arabidopsis. PLOS Pathogens 9: e1003370.

Zhu M, Assmann SM. 2017. Metabolic Signatures in Response to Abscisic Acid (ABA) Treatment in Brassica napus Guard Cells Revealed by Metabolomics. Scientific Reports 7: 12875.

Zilberman D, Cao X, Johansen LK, Xie Z, Carrington JC, Jacobsen SE. 2004. Role of Arabidopsis ARGONAUTE4 in RNA-Directed DNA Methylation Triggered by Inverted Repeats. Current Biology 14: 1214–1220.

